# Validation of diverse and previously untraceable Sendai virus copyback viral genomes by Direct RNA Sequencing

**DOI:** 10.1101/2025.04.04.647164

**Authors:** Sarah E. Pye, Emna Achouri, Yanling Yang, Abdulafiz Musa, Carolina B. López

## Abstract

Copyback viral genomes (cbVGs) are truncated viral genomes with complementary ends produced when the viral negative-sense RNA virus polymerase detaches from the replication template and resumes elongation from the nascent strand. Despite advances in methods to identify cbVGs based on the site of polymerase break and rejoin, PCR-based tools cannot provide full length sequences of most cbVGs and/or can introduce errors and artifacts during cbVG amplification. These limitations have painted a limited picture of the diverse population of cbVGs generated during infection. To improve our ability to obtain native full-length sequences of cbVGs, we optimized Direct RNA Sequencing (DRS) as a fast and simple tool to sequence full-length cbVGs and designed a BLAST-based analysis approach to identify cbVGs from long-read sequencing data. We analyzed the DRS outputs of multiple Sendai virus stocks to highlight both the utility and limitations of this tool. We found that to capture the dominant 546 nucleotide cbVG produced by Sendai virus strain Cantell, the length of complementarity between the virus trailer and the DRS oligonucleotide should optimally be increased to up to 32 nucleotides. We also demonstrate comparable quality of cbVG sequences by DRS from as little RNA as 17.6ng from the media fraction or 50ng of from the cellular fraction of cells infected with SeV, in contrast to the recommended 1000ng. Importantly, we validated different cbVG species from two recombinant Sendai virus stocks, including for the first time cbVGs whose break positions occurred at or near position one in the reference genome.

**Importance:** Most viruses of the order Mononegavirales has been demonstrated to naturally generate copyback viral genomes. These genomes are critical determinants of infection outcomes; they interfere with standard virus replication by competing for viral resources, activate antiviral responses, and inhibit protein translation. Despite their critical roles in infection, current tools to study copyback viral genomes rely either on preexisting knowledge of the sequence of a target RNA or require reverse-transcription and amplification of the target RNA, biasing toward short cbVGs and introducing relatively high rates of errors. Our lab has long advocated for RNA virologists to sequence their stocks to assess the role cbVGs may have in their infections. Toward this effort, we detail the optimization of Direct RNA Sequencing to sequence full-length cbVGs while maintaining the integrity of the native cbVG RNA and present the Long-Read cbVG Analysis (LoCA; https://github.com/lopezlab-washu/LOCA.git) script as a highly accessible BLAST-based tool for cbVG detection.

## Introduction

Negative-sense (ns)RNA viruses generate a diverse population of viral particles defined by their genomic content. Viral genomes include the standard viral genomes copied exactly from the positive sense antigenomic intermediate, and replication-competent viral variants that harbor non-lethal mutations (1). RNA viruses also produce nonstandard viral genomes (nsVGs), whose replication requires the standard virus polymerase (2, 3). In nsRNA viruses, the majority of these nsVGs are copyback viral genomes (cbVGs) generated when the viral polymerase detaches from the replication template at a “break position” and resumes elongation downstream at a “rejoin position”, copying back a sequence complementary to the 5’-end of the nascent genome (Fig. 1A, adapted from (4)) (1). cbVG generation appears to be restricted to but conserved among negative-sense RNA viruses, with cbVGs identified within every virus of the order Mononegavirales (1, 5). To understand how RNA viruses establish infections, promote transmission to other hosts, and are maintained within the host population, it is important to consider all components of the virus community, including both standard and nonstandard viral genomes.

**Figure 1:**
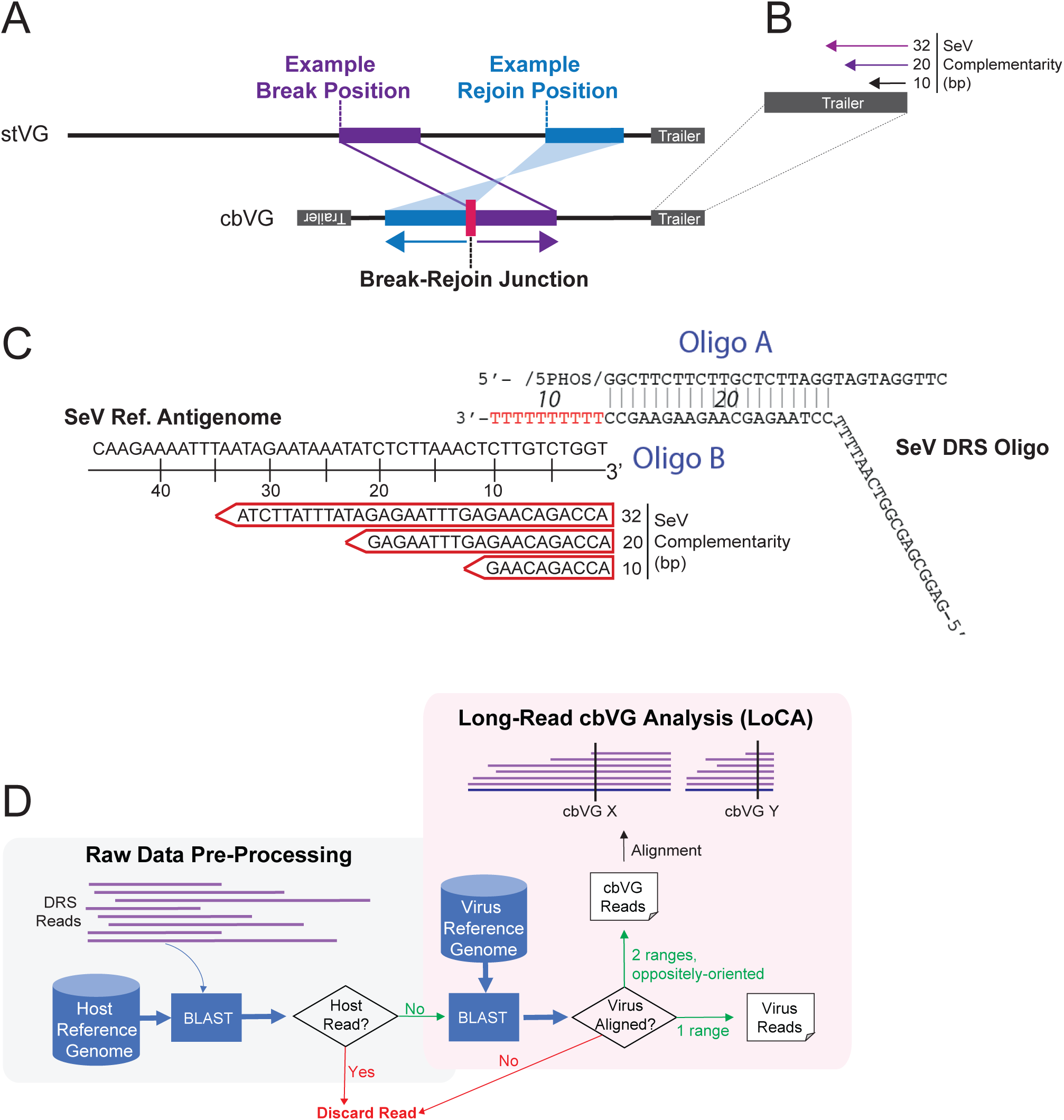
**LoCA identifies copyback viral genomes (cbVGs) from long-read Direct RNA sequencing data**. (A) cbVGs are generated when the viral polymerase detaches from the replication template at a “break position” and resumes elongation downstream at a “rejoin position”, copying back a sequence complementary to the 5’-end of the nascent genome. Adapted from [22]. (B) Sequence-specific adapter oligonucleotides of lengths 10, 20, and 32 nucleotides were designed and tested against the Sendai virus (SeV) trailer for Direct RNA Sequencing (DRS). Oligo design in trailer permits cbVG capture independent of break-rejoin junction. (C) The final DRS adapter oligonucleotide is annealed by mixing Oligo A and Oligo B 1:1 in buffer (1.4 μM in 10 mM Tris-HCl pH 7.5, 50 mM NaCl), heating to 95° C for 2 minutes, and cooling slowly at 0.1° C/sec. Oligo A is an ONT-specified sequence while Oligo B contains 10 dT nt, which were replaced by the corresponding SeV binding sites represented in red arrows. (D) The DRS workflow for cbVGs begins with the removal of host-aligned reads from the sequencing data. Next, NonHost reads are aligned by BLASTn to the virus reference genome from which reads with one alignment range are reported as virus reads and reads with two alignment ranges in opposite orientations are reported as cbVG reads. cbVG reads with common break-rejoin junctions are grouped into a single cbVG species.

During infection, cbVGs interfere with standard virus replication by competing for viral resources, activating antiviral responses, and by inhibiting protein translation (6). We have demonstrated that cbVGs arise naturally *in vivo* during infection with respiratory syncytial virus (RSV) in children and with Sendai virus (SeV) in mice (7–9). cbVGs were established as the primary triggers of robust antiviral responses to RSV in children, making them candidate biomarkers to predict the outcome of infections and the rate of virus spread in a population (8). Timing matters for this effect; detection of cbVGs early after infection is associated with low viral load and mild disease, where worsened clinical outcomes are seen in patients that generate cbVGs late in infection or present for prolonged periods (10). With mounting evidence that nsVGs are naturally produced and amplified during infection, cbVGs are now proposed as agents for therapeutic interventions (11–14).

To investigate the impact of specific cbVGs in an infection, one must first know which cbVGs are present. Existing strategies typically employ Illumina Sequencing to predict cbVG species complemented by RT-PCR to validate them. cbVG detection from short read sequencing data is based on the reference genome sequence, where cbVG reads contain a junction of two oppositely oriented regions of genome sequence, called a break-rejoin junction or position (Fig. 1A). While Illumina sequencing is highly sensitive and has dramatically increased our ability to detect cbVGs from diverse samples, the requisite fragmentation and short read length are severely limiting when trying to study full-length cbVGs, especially the presence or function of their molecular motifs. Further, like all RT-PCR based methods, Illumina sequencing relies on cDNA synthesis which can create artificial chimeric cDNA from template switching during RT and mis-priming in PCR (15–17). Thus, cbVG research can be limited by artifacts introduced both during the initial identification of a cbVG from Illumina sequencing and in its validation by RT-PCR.

By contrast, Oxford Nanopore Technologies’ (ONT) Direct RNA Sequencing (DRS) eliminates biases from reverse transcription and/or amplification and does not require fragmentation of the RNA, allowing for sequencing of full-length native RNA molecules. In DRS, native RNA molecules pass through protein nanopores embedded in an electrically charged membrane, with each nucleotide characteristically disrupting the ionic current to generate sequencing reads. DRS has already been appreciated for its ability to reveal unexpected transcriptional and genomic complexity from viral pathogens (18–20).

To optimize DRS for sequencing of full-length cbVGs, we utilized SeV because it is a well-studied murine paramyxovirus related to human parainfluenza viruses (7, 21–23). Further, we employed the historically available Sendai virus strain Cantell because it naturally produces and accumulates one specific well-characterized cbVG of 546 nucleotides (nt) long (cbVG 546), that confers strong immunostimulatory ability to the virus and shuts off protein translation (7, 24–27). We detail the optimization of DRS for sequencing full-length cbVGs and apply these findings to gain novel insight into the cbVG populations of two virus stocks with diverse cbVG species. While DRS was limited in its ability to quantitative absolute cbVG numbers, the tool successfully validated full and partial-length cbVGs of variable lengths, including for the first time cbVGs with break and rejoin positions at opposite ends of the reference genome, in accordance with our previous predictions from both, virus passaged in vitro and clinical samples (28). Apart from some partial length cbVGs whose 5’ read ends terminated prematurely, all the cbVGs we validated were sequenced in their entirety, with no evidence of internal deletions (also called “mosaic” cbVGs). We show that DRS is a fast and accurate tool for validation and characterization of cbVGs from virus stocks and infected samples.

## Results

### DRS optimization for cbVG detection

Direct RNA Sequencing kits sold by ONT include an adapter oligonucleotide that contains 10 d(T) nt, designed to bind the terminal 3’ end of mRNA, within the poly A tail. To capture cbVGs originating from the trailer, we designed our adapter oligonucleotide to bind the terminal nucleotides in the trailer sequence of the SeV antigenome. This positive-sense trailer sequence is present in all cbVGs regardless of their specific break-rejoin junction, as well as in the standard full-length viral antigenomes (Fig. 1A). However, we noted that adapter oligonucleotides designed to target the terminal 10nt of the antigenome would have several potential off-target binding sites within the SeV genome due to similarity in sequences. To investigate whether the adapter oligonucleotide could be optimized by increasing the oligonucleotide length, we designed and tested DRS adapter oligonucleotides of 10, 20, and 32 nt complementarity (Fig. 1B-C). Initial optimization experiments were performed at the manufacturer’s recommendation of 1000ng input (total) RNA, collected from SeV-infected A549 cells 24 hours post infection (hpi) at a multiplicity of infection (MOI) 1.5 TCID_50_/cell. Data were analyzed using our publicly available Long-Read cbVG Analysis script (LoCA; https://github.com/lopezlab-washu/LOCA.git). Host read filtered sequencing data can be input into LoCA, which only requires a BLASTn nucleotide database for the virus reference genome. In summary, LoCA sorts and reports reads that align in exactly one range as virus reads and reads that align in two oppositely oriented ranges as cbVG reads (Fig. 1D).

First, we investigated the requisite length of virus complementarity within the adapter oligonucleotides for cbVG capture and indeed found that the 10nt long adapter oligonucleotide was suboptimal to sequence cbVG546. By contrast, longer adapter oligonucleotides of up to 32nt increased each the total number of non-host (total) reads (Fig. 2A), virus reads (Fig. 2B), and cbVG reads (Fig. 2C). We next investigated whether increasing the concentration of the adapter oligonucleotide from the recommended 2μM to 4μM or 8μM would increase the number of cbVGs and stVGs but found that providing more adapter oligonucleotide decreased cbVG capture (Fig. 2D). In addition, excess adapter oligonucleotides during library preparation increased the frequency of background reads (non-host and non-virus aligned) (Fig. 2E). Further, as expected when sequencing a stock that makes one short dominant cbVG, most reads in a representative sequencing experiment were shorter than 2kB, but this strategy also captured some nearly full-length (15kB) standard viral genome reads (Fig. 2F).

**Figure 2:**
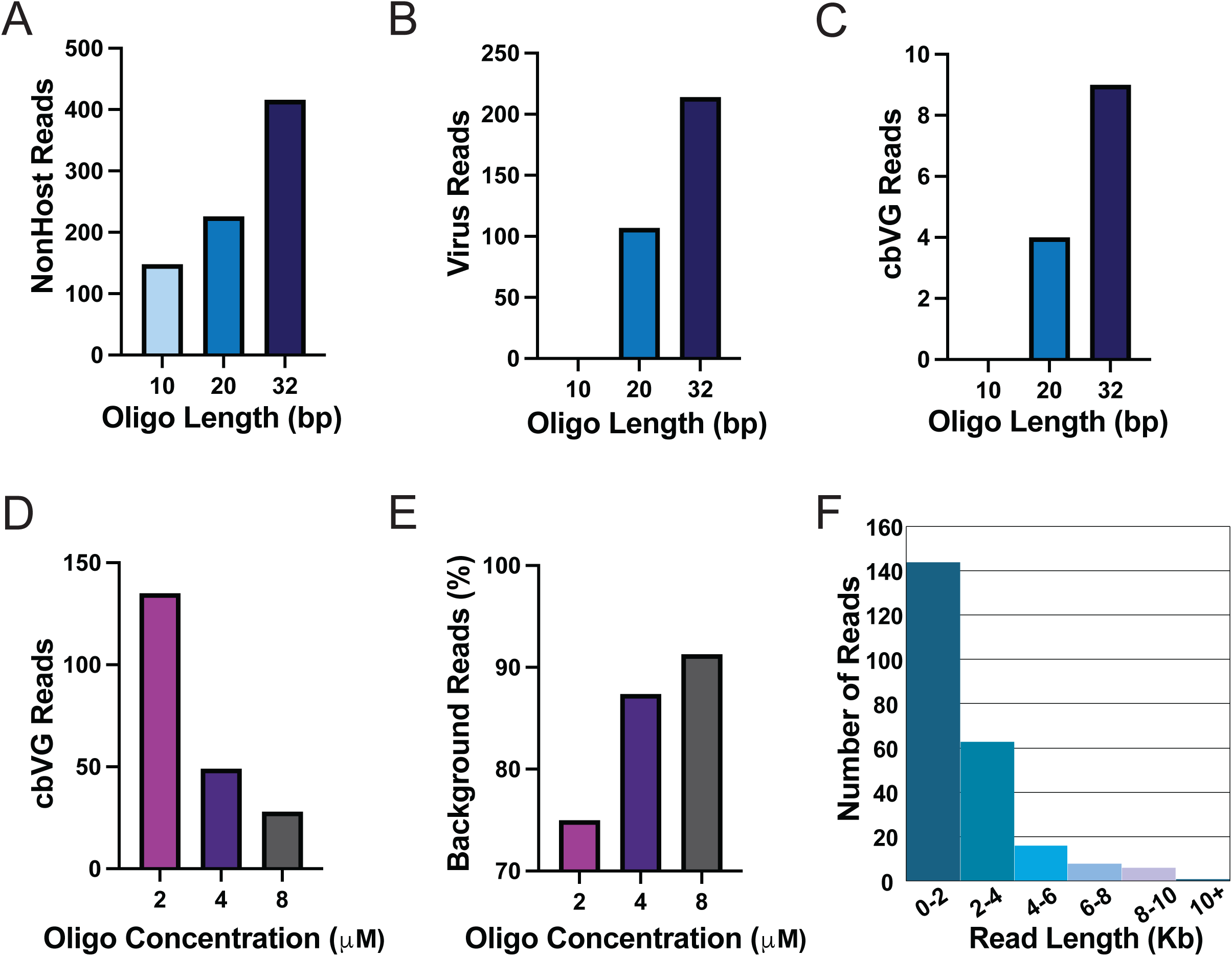
The length of sequence-specific oligonucleotide is critical for capture of cbVGs. DRS output from cellular RNA extracted from A549 cells 24 hours after infection by SeV Cantell at MOI 1.5. The library was prepared using sequence-specific adapter oligonucleotides with the indicated lengths of complementarity to the virus trailer and sequenced for 72 hours. Increasing the length of trailer-specific adapter oligonucleotide improved capture of (A) total (non-host) reads, (B) virus-aligned reads, and (C) cbVG reads. However, increasing the adapter oligonucleotide concentration from 2μM to 8μM decreases capture of (D) cbVG reads and increases (E) background reads. (F) Histogram showing the lengths of Sendai virus-aligned reads in an example DRS sample using 2μM of a 32nt trailer-specific adapter oligonucleotide. Read length bins every 2,000 nucleotides.

### DRS application along a time course

Next, we validated DRS in SeV infections of A549 cells over a time course, collecting both cellular fraction RNA (to capture active replication in the cells) and the more dilute media fraction RNA to reduce contamination by host RNA. In panels A and B of Figure 3 are two independent experiments sequencing cellular RNA 6 - 24hpi. In panel C is one experiment sequencing RNA from the media fraction 12-48hpi. We found that 6hpi is too early to capture cbVGs by DRS, that cbVGs are detectible although more variable between experiments at 12hpi, and that cbVG levels are consistently sufficient and comparable between experiments beginning at 24hpi (Fig. 3 A-C). A detailed alignment of the DRS generated cbVG 546 sequence from the 24hpi, 1000ng cellular RNA condition in Panel A of Fig. 3 can be found in Fig. S1. The cbVG 546 sequences are high quality, with mismatches typically occurring as deletions within nucleotide repeats, also called homopolymers (Fig. S1). This type of error is expected because cbVGs are highly structured and have shorter read lengths, resulting in more variable translocation speeds than stVG. When sequencing long stretches of the same base by DRS, the signal does not change, so if the RNA translocation speed is not constant, the number of bases in homopolymeric regions can be misrepresented (29).

**Figure 3:**
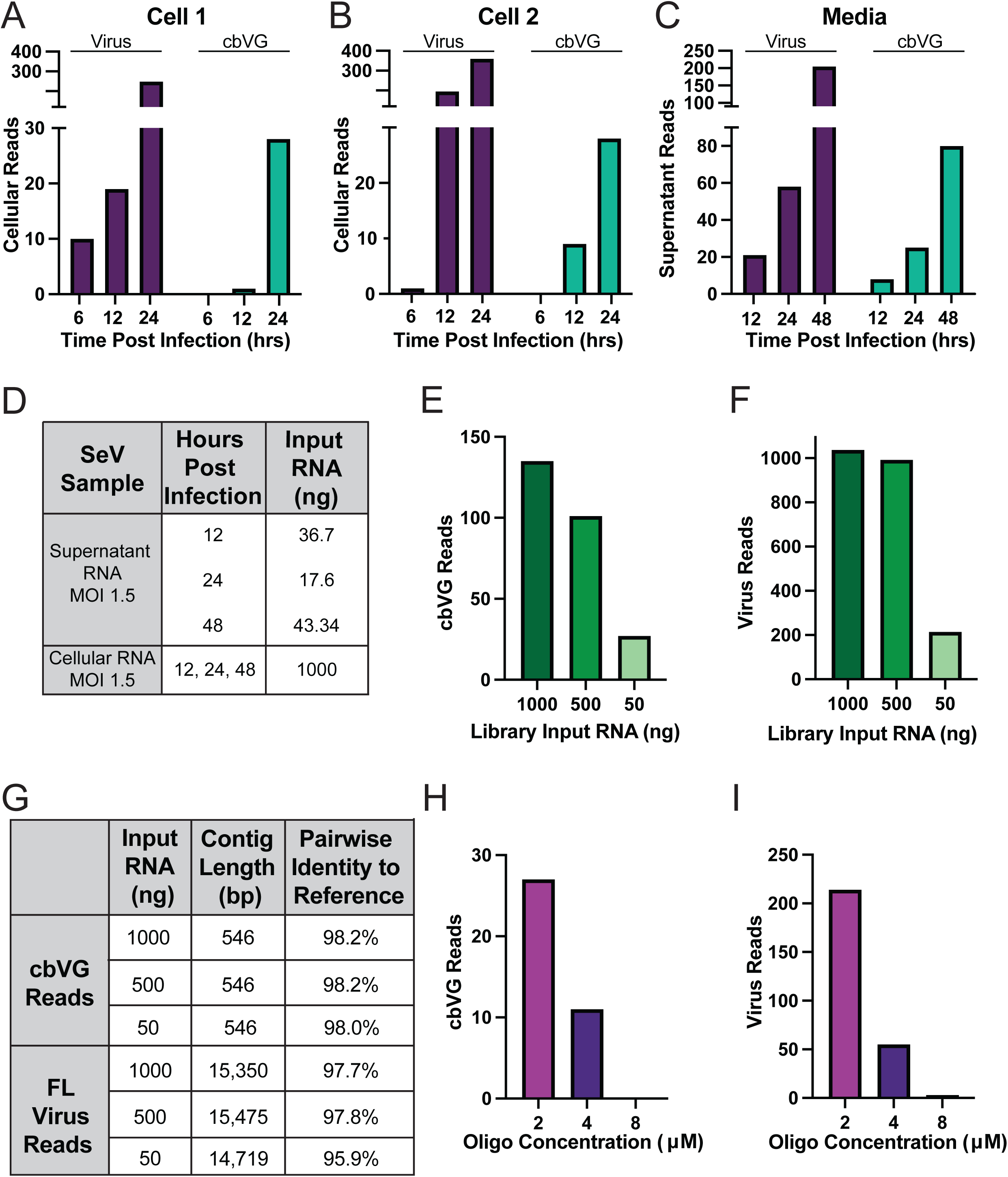
DRS can be used to sequence SeV copyback and standard viral genomes from RNA extracted from both cells and media. Infections of A549 cells were performed using Sendai virus strain Cantell at MOI 1.5 for indicated time points. Libraries were prepared using sequence-specific adapter oligonucleotides with 32nt complementarity to the virus trailer and sequenced for 72 hours. (A-C) Virus and cbVG reads by sequencing libraries from either the cellular or media RNA fractions as indicated. (D) Library input RNA for each condition of the cellular and media time course experiments in Panels A-C. (E-F) cbVG and virus reads identified from cellular fraction RNA libraries at indicated input RNA quantities. (G) Percent identity and lengths of resulting cbVG and virus sequences for each condition of the cellular and media time course experiments in Panels E-F. (H-I) cbVG and virus reads from varying the concentration of trailer-specific adapter oligonucleotide from 2μM to 8μM while maintaining 50ng library input RNA.

Of note, while all cellular RNA experiments sequenced a library made from 1000ng input RNA, far less than 1000ng was extracted from the media fractions (Fig. 3D). However, DRS still produced high quality complete cbVG546 sequences from as little as 17.6ng of input RNA (Fig. 3D). Since the media fraction is more diluted and has less host RNA than the cellular fraction, we wondered if cbVG sequencing will still be successful after decreasing the input RNA from the cellular fraction, where cbVGs are selected for among a greater population of potential contaminants. To test this, we decreased the cellular fraction input RNA to 500ng and 50ng and found reduced numbers of both cbVG and virus reads (Fig. 3E and 3F). However, pairwise identity was comparable between the cbVG sequences obtained from the 50ng and 1000ng conditions (Fig. 3G). These results suggest that for most applications, 50ng of total RNA is sufficient for sequencing cbVGs. For assessments of pairwise identity, the DRS sequence was aligned to a hypothetical full-length reference cbVG 546 sequence generated to contain every nucleotide from the end of the trailer until the break position, immediately followed by every nucleotide from the rejoin position back to the end of the trailer. As before, offering more adapter oligonucleotide during library preparation did not improve capture of cbVG or full-length virus (Fig. 3H and 3I).

### DRS validates diverse cbVGs predicted from Illumina data

We next evaluated whether DRS can validate and provide complete cbVG sequences in a system with more diverse cbVGs that includes longer species not previously validated. To do this, we employed two newly generated high cbVG producing stocks of rescued recombinant SeV, named rSeVA and rSeVB, which have cbVG profiles distinct both from each other and from the cbVG 546-producting historical SeV Cantell stock described above. First, we used previously designed primers (7) to detect cbVGs in the rSeVA and rSeVB stocks by PCR, producing the bands in Fig. 4A. Whole plasmid sequencing after these bands were cloned into pGEM-T plasmids validated the break-rejoin junctions of four short cbVG species as listed in Fig. 4B. Of note, these are partial sequences, limited by the location of the primers. Further, the enzymatic nature of RT-PCR also biases shorter species, which can be limiting when validating very long cbVGs. For these and many other viruses, RT-PCR for cbVG validation is extremely low throughput as compared to Illumina sequencing.

**Figure 4:**
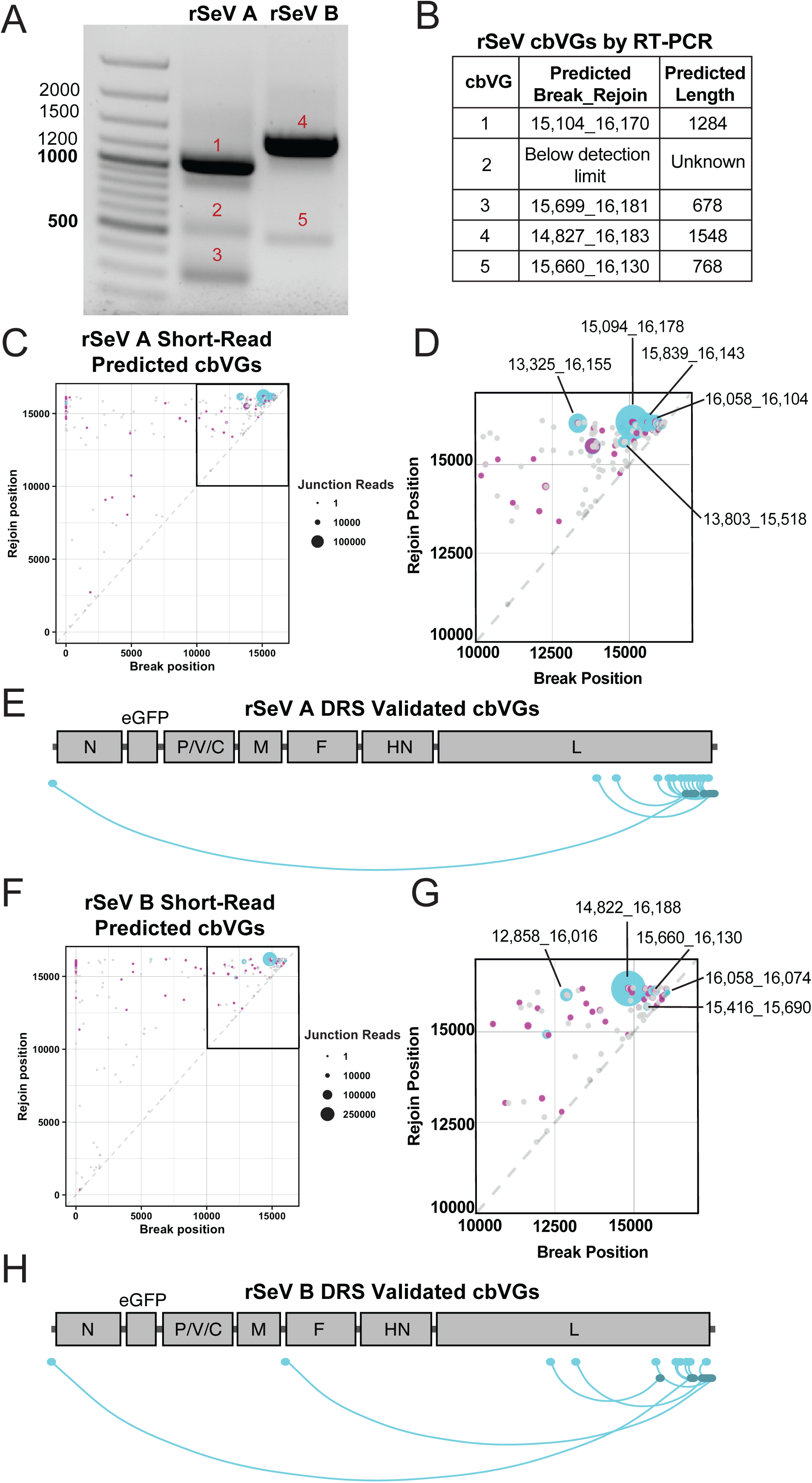
DRS validates cbVGs from rSeV A and B stocks. (A) Agarose gel (1%) highlights amplicons resulting from cbVG-specific RT-PCR of rSeVA (left) and rSeVB (right) virus stocks. Bands from the gel were excised, purified, and sequenced. (B) Break-rejoin junctions and predicted full lengths of sequence-confirmed RT-PCR validated cbVGs. Numbering of break-rejoin junctions and predicted lengths of cbVGs assume synthesis of every nucleotide from the trailer until the break, and from the rejoin back to the end of the trailer of the SeV reference genome. (C, F) Total population of cbVGs predicted by VODKA2 from Illumina short read sequences of rSeVA and B stocks respectively (magenta dots). cbVG break-rejoin junctions validated by DRS designated as cyan dots. (D, G) Zoom view for rSeV A and B stocks respectively, highlighting the high frequency of predicted and validated cbVGs characterized by break-rejoin junctions nearest the virus trailer. (E, H) Virus gene annotations for break and rejoin positions of cbVGs validated by DRS at three or more reads from rSeV A and B stocks respectively. Break position is indicated in cyan, joined to the corresponding rejoin position in navy.

Thus, we next used Illumina sequencing to generate short sequencing reads from these samples, which we analyzed by VODKA2 to predict cbVGs from each rSeVA and rSeVB stocks (30). Briefly, VODKA2 uses Bowtie2 to align the short reads from Illumina sequencing to a catalog of hypothetical break-rejoin junctions, generated based on the sequence of the reference genome (30). VODKA2 then validates by BLASTn that the predicted cbVG read is aligned in two oppositely oriented ranges, before generating a report of all predicted cbVGs (30). By contrast to RT-PCR, analysis of Illumina sequencing data by VODKA2 detects hundreds of thousands of cbVG break-rejoin junctions. Reads with similar break-rejoin junctions and lengths can be classified as cbVG species. We represented each cbVG species as a magenta dot whose size is correlated with its predicted frequency on a scatter plot, whose axes represent the position of each cbVG break (x-axis) and rejoin (y-axis) in the reference genome (Fig. 4 C, F). However, these break-rejoin junctions are only predicted and must undergo secondary validation like RT-PCR or DRS.

Thus, to validate and obtain complete sequences of the more complex cbVG population predicted from short read data, we sequenced the rSeVA stock twice and the rSeVB once by DRS, each time with 50ng of RNA as input for the library. DRS validated 16 predicted cbVGs with N≥3 reads from rSeVA and 9 predicted cbVGs with N≥3 reads from rSeVB. DRS validated cbVG species (N≥3) DRS reads with shared break-rejoin positions ± 5nt are indicated as cyan dots in Fig. 4C and 4F for rSeVA and rSeVB respectively. Full-length DRS-validated cbVGs ranged in length from 396nt to 3240nt. These included both cbVG species validated by RT-PCR from each stock in Fig. 4 A-B.

Of note all cbVGs sequenced at N≥3 DRS reads followed the paramyxovirus “rule of 6” where the paramyxovirus genome size must be a precise multiple of six to be replicated efficiently (Fig. S2, S3). As expected, and as reflected by the increased number of DRS reads and corresponding larger dots on the scatter plot, most of the validated cbVG break-rejoin junctions are near the trailer end of the genome (zoomed views in Fig. 4D, 4G). For the first time, this tool also enabled us to validate cbVG break-rejoin junctions in each sample characterized by distal break positions as far as position 1 in the reference genome, with rejoin positions in or near the trailer (Fig 4E, 4H). Of note, although not every Illumina cbVG was validated, all cbVGs sequenced by DRS corresponded to a cbVG predicted from the Illumina data. A complete list of all validated cbVGs and their properties is provided in Fig. S2 for rSeVA and Fig. S3 for rSeVB.

### DRS cbVG reads are high quality

During basecalling of DRS data, each read is assigned a quality score (Q score), frequently used as a filter to exclude poorly sequenced and basecalled reads (31). Q10 corresponds to an error rate of 10% and Q20 corresponds to an error rate of 1% (31). The default recommended DRS Q score threshold is Q=9, by which we find that the cbVG reads are, on average, very high quality. Specifically, Fig. 5A reports an average score of 21 from a fastQC analysis conducted on the pooled rSeVA and rSeVB DRS cbVG reads basecalled at Q=9, corresponding to an average basecall accuracy of 99.2%. However, the Q score threshold is a parameter that can be modified by the user to include more prospective sequences if desired. When we decrease from Q=9 to Q=7, total cbVG read numbers increase 69.7%, with some cbVG species increasing dramatically more than others (Fig. 5B). We find that many of the cbVG reads with Q scores <9 are partial length cbVGs, so while these are not our target, they may prove useful to a user searching for the maximum possible number of reads corresponding to a specific cbVG.

**Figure 5:**
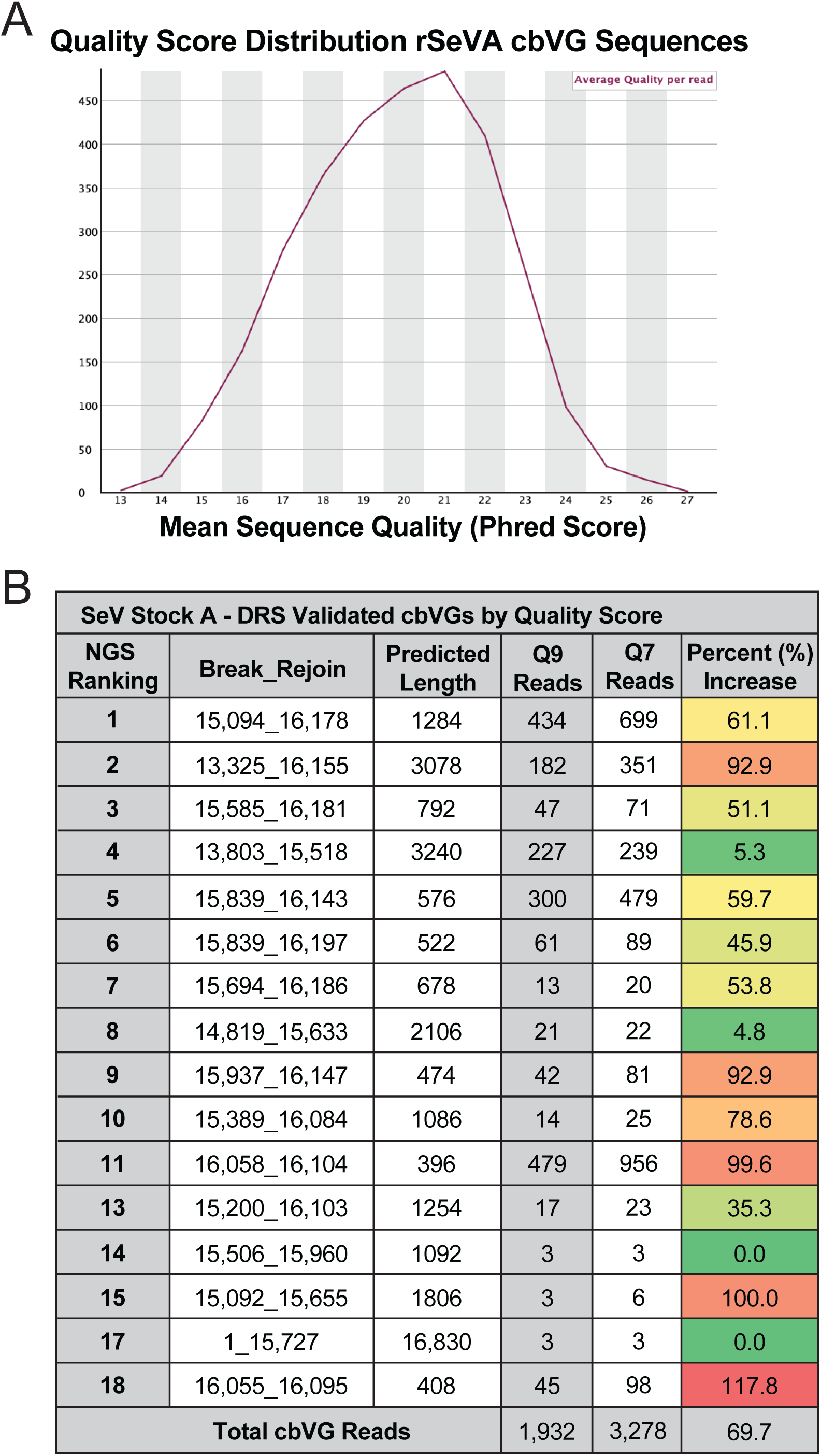
cbVGs sequencing reads from DRS are high quality. (A) FastQC report showing quality per read of all DRS validated cbVG reads from both rSeVA and rSeVB stocks. Each sample basecalled at a minimum quality score threshold of 9 prior to fastQC analysis. (B) The quality score (Q-score) threshold can be modified by the user during basecalling and maintains or increases DRS read numbers for each break-rejoin junction species. Chart shows the NGS ranking corresponding to the relative frequency of detection of each break-rejoin junction by NGS, the predicted length, the number of DRS reads at the recommended quality score 9, and the number of DRS reads after reducing the quality score threshold to 7. The final column shows the percent increase in read counts by decreasing the quality score threshold to 7, colored as a heat map with the smallest increases in green and the largest increases in red.

### DRS validates distal break-rejoin junctions

Diagramed in Fig. 6A is the longest full-length cbVG sequenced, cbVG3240 (from rSeVA; named for its full length in nt). In general, cbVG sequence integrity by DRS is high; across all experiments, mismatches observed by DRS were typically deletions instead of substitutions. This holds true for cbVG 3240, where all 20 mismatches were deletions, often in regions of repeated nucleotides reflecting a technical limitation of DRS, not reflecting a biological feature of cbVGs (Fig. 6B). Concerning the overall architecture of full-length cbVGs, we did not observe any large internal deletions or rearrangements resembling mosaic cbVG architecture.

**Figure 6:**
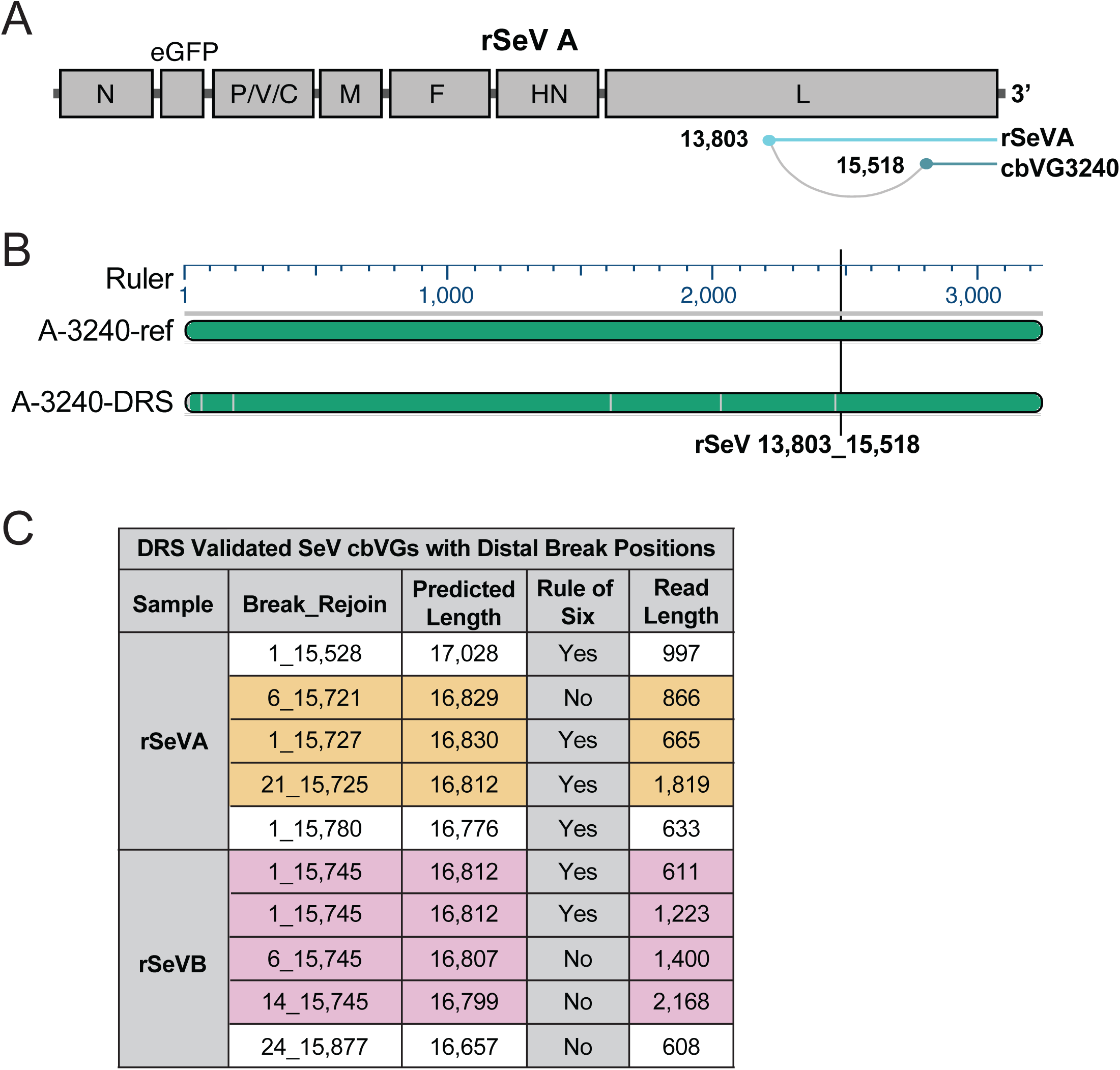
DRS validates the existence of long cbVGs by their break-rejoin junctions. (A) Illustration of the architecture of the longest full-length cbVG validated by DRS, rSeVA-cbVG 3240 (named for its predicted length). Sequence until cbVG break at rSeV position 13,803 represented in cyan, with sequence from cbVG rejoin at rSeV position 15,518 until the end of the trailer represented in navy. (B) Alignment overview of rSeVA-cbVG 3240, highlighting that the full length of the predicted cbVG is sequenced; there was no evidence of mosaic cbVG architecture characterized by internal deletions. The reference cbVG sequence is on top with the DRS generated sequence on bottom. Grey vertical lines in DRS sequence indicate deletions from reference sequence (no substitutions or insertions were present). (C) DRS reads validate cbVGs with distal break positions at or near position 1 in the reference genome. Shown are the predicted complete length and actual read length of each cbVG with its validated break-rejoin junction, and whether it follows the paramyxovirus “rule of six” (genome length = 6n+0). cbVG species are highlighted from each sample, grouped by shared rejoin position of +/-6nt.

Finally, as mentioned previously, these DRS experiments validated cbVGs in both rSeV stocks characterized by a distal break at position 1 in the reference genome, and a rejoin in the final gene (L, polymerase gene) encoded by the virus genome. Of note, some of these reads also follow the rule of 6, as highlighted in Fig. 6C. This is the first direct validation that these species exist, which have long been predicted among clinical and complex infection samples (10, 28). Like the diagram in Fig. 6A, the length from the trailer to the break position in these cbVGs is much longer than the length from the rejoin position back to the trailer, where the oligonucleotide is positioned. This enables us to directly validate for the first time break-rejoin junctions from cbVGs with predicted full lengths up to 16,830nt, even though we don’t sequence every nucleotide. Previously, we have lacked adequate tools to validate whether these are true cbVGs or simply artifacts introduced during the library preparation for Illumina sequencing. By DRS, we are highly confident in the verity of these cbVG species.

### DRS shortcomings

DRS would be maximally helpful to us as a quantitative tool, to assess how many cbVGs are produced compared to standard viral genomes, or to compared cbVG accumulation levels between species. However, even in the most optimal conditions tested, we found that a large percentage of the reads aligned to the host genome or were called as background reads. To dive deeper into this issue, we investigated which RNAs were represented in the reads from the 8μM oligonucleotide condition, where nearly 80% of all sequenced reads align to the host genome (Fig. S4A). Among the 20% non-host reads, most reads aligned to the standard virus, with 7.4% cbVG reads and 29.4% background reads (non-host and non-virus aligned) (Fig. S4A). The host reads were enriched in the most prominent cellular RNAs in a typical A549 cell. Specifically, chromosome 17 is the most highly sequenced because of a polysomy of this chromosome in A549 cells (32), followed by ribosomal RNAs, long non-coding RNAs, and mitochondrial RNAs (Fig. S4B). Examination of the background reads in this sample showed that most cannot be classified as they are primarily composed of either G/C or A/U repeats (Fig. S4C). The background reads that could be manually classified included some very short reads aligning to common host RNAs (Fig. S4C). The high level of host reads sequenced from infections in A549 cells was not seen from infections in highly permissive monkey LLCMK2 cells (Fig. S4D).

To reduce the proportion of host reads sequenced from A549-infected cells, we tested two methods for host RNA depletion before library preparation: Ribozero ribodepletion and Dynabeads mRNA depletion, as well as adaptive depletion through MinKNOW during sequencing for targeted depletion of the most common host reads sequenced previously. However, all these strategies similarly failed to significantly reduce the percentage of host reads in the sample (Fig. 7A). To verify successful ribodepletion, we confirmed by Bioanalyzer that the 18S and 28S peaks present in total RNA infected with SeV (Fig. 7B) are depleted after Ribozero Ribodepletion (Fig. 7C), which can be confirmed using a higher sensitivity assay (Fig. 7D). Comparing Ribozero Ribodepletion before library preparation to Adaptive Depletion during MinKNOW during data acquisition, we did not find proportionally higher levels of virus or cbVG reads, although we did sequence more reads overall compared to DRS without any host depletion, with increases in total numbers of both virus and cbVG reads (Fig. 7E).

**Figure 7:**
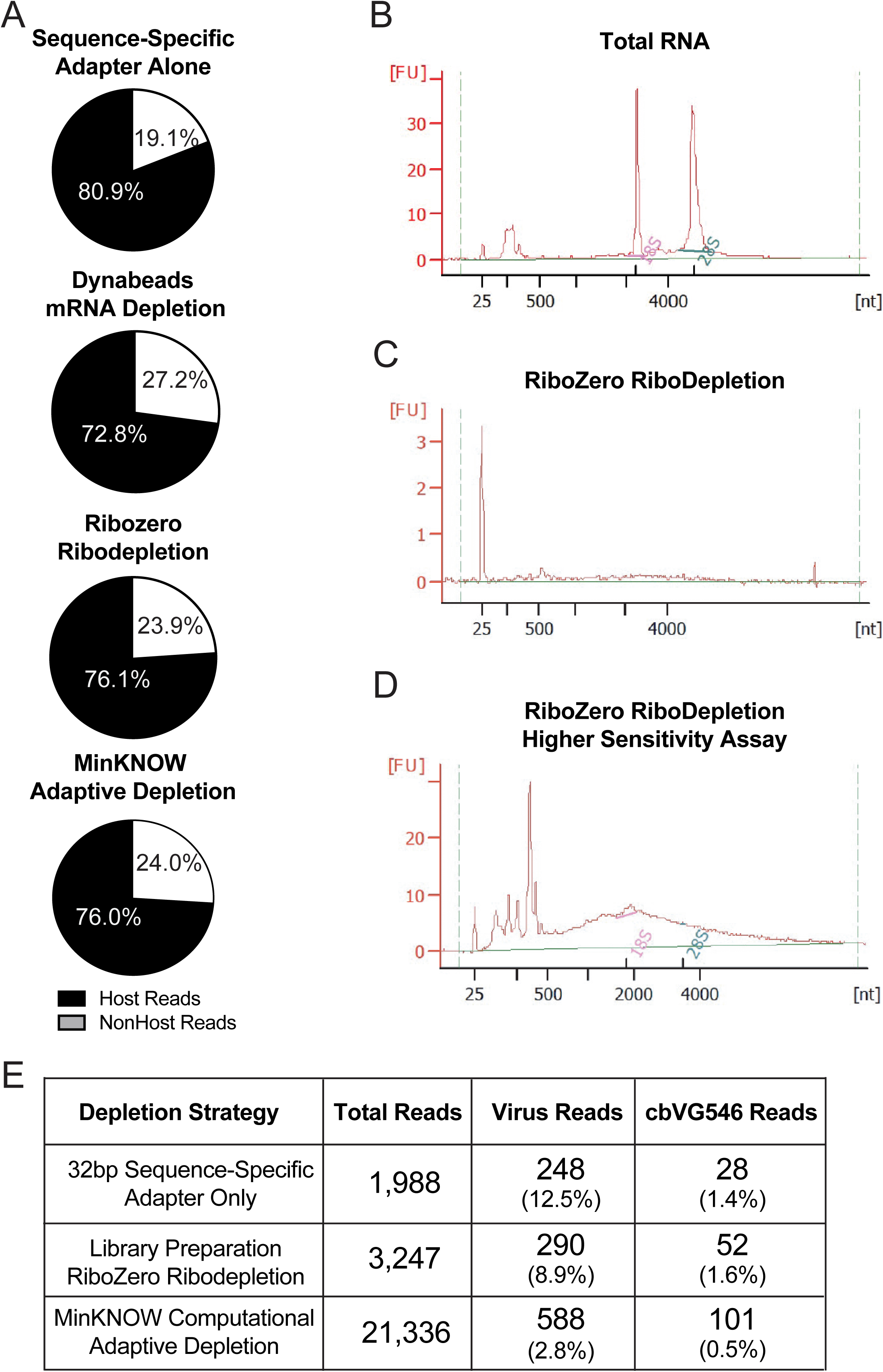
DRS in A549 cells is limited by nonspecific capture of host reads. (A) Percentages of host reads (black) and non-host reads (white) from sequencing DRS libraries prepared using the sequence-specific adapter alone, in addition to Dynabeads mRNA depletion or Ribozero ribodepletion, or from MinKNOW adaptive depletion computationally during data acquisition. (B) Bioanalyzer electropherogram traces from total RNA with 18S and 28S peaks labeled in pink and green respectively. (C) Bioanalyzer electropherogram traces from Ribozero ribodepleted RNA. (D) Higher sensitivity bioanalyzer electropherogram traces from Ribozero ribodepleted RNA with 18S and 28S peaks labeled in pink and green respectively. (E) Comparison of the number of total reads, virus-aligned reads, and cbVG reads from DRS of total RNA using the sequence-specific adapter alone, Ribozero ribodepleted RNA, and the sequence-specific adapter followed by MinKNOW adaptive depletion computationally during data acquisition. The percentages of virus and cbVG 546 reads are shown in parentheses below the read numbers.

## Discussion

As cbVG generation is broadly conserved among the Mononegavirales, and cbVGs are highly influential in infections, it is critical that virologists have quick, simple, and reliable tools to identify cbVGs from virus stocks and infection samples. Toward this end, we describe the optimization of DRS to sequence full-length cbVGs and its application to simultaneously sequence multiple variable length cbVGs at the native RNA level without potential amplification-induced errors. We first found that to prepare successful libraries for DRS of cbVGs, users could increase the length of SeV sequence-specific adapter oligonucleotides complementary up to 32 nt. We also found that preparing libraries with only 50ng of input RNA is sufficient for most applications. Further, we showed that depletion of host RNA, either by Ribozero Ribodepletion or by MinKNOW Adaptive Depletion, increased the total number of sequenced reads and the number of both virus and cbVG reads, although didn’t increase the percentage of virus or cbVG reads in the sample. In these cases, removing host reads from the sample that could otherwise occupy and clog nanopores enabled the sequencing of more RNA molecules in total, although this did not affect the proportion of host RNA likely because the adapter oligonucleotide binds nonspecifically to prominent RNAs in the host cell. Together, this suggests that DRS is sufficient to sequence full-length cbVGs and stVG but is not yet quantitative.

Furthermore, we describe LoCA (https://github.com/lopezlab-washu/LOCA.git), an entirely BLAST-based analysis workflow to identify cbVGs (virus aligned in two orientations) from long-read sequencing data. Existing cbVG detection pipelines are specialized for Illumina reads and classify cbVG reads by aligning the experimental data to a catalog of hypothetical break-rejoin rejoins (30). The BLAST-based analysis we describe here to identify cbVGs from long-read sequencing data offers several advantages by comparison. First, there is reduced computational and labor demand because it is only necessary to generate and store a BLAST nucleotide database for the reference genome, not a catalog of hypothetical break and rejoin positions. This strategy is also unbiased and highly specific because it classifies cbVG reads based on alignment ranges instead of percent identity to a predicted reference. Finally, this also enables classification of partial or mutated reads; cbVGs can be reported independent of their predicted or actual length, and regardless of whether there are mutations or even truncations outside of the break-rejoin junction. In all the optimization and application experiments, cbVG reads reported by LoCA were true cbVG reads; there was no evidence of artifactual recombination like we have to consider when using other methods. One limitation of this study is that we analyzed LoCA reads with precisely two alignment ranges; we may have detected cbVGs with more complex architecture by investigating reads with more than two alignment ranges.

When we applied DRS to SeV stocks with more diverse populations of cbVGs, we validated 16 cbVGs from two sequencing experiments and 9 cbVGs from one sequencing experiment, each using only 50ng RNA per library. Full-length (>99%) validated cbVGs ranged in length from 396-3240nt and in the case of partial length cbVGs sequenced by DRS, we find that the 5’ end of the cbVG sequence may be incomplete. As in DRS RNA molecules are translocated across the pore 3’->5’ but flipped during basecalling to generate 5’->3’ oriented reads, it is expected that the 5’ ends of some reads are missing as the most distal sequences supposed to pass to the pore can be lost due to RNA degradation or nanopore clogging. Among these partial length cbVGs, we validated for the first time the existence of cbVGs with distal break-rejoin junctions, specifically those with break positions at/near position 1 in the reference genome and rejoin positions in the trailer region. These very long cbVGs may be an intermediate species during the diversification of cbVG species during infection (28). We hypothesize that long cbVGs occur early during infection an act as templates for smaller cbVGs generated later during infection, One limitation of DRS is that cbVGs with a very short loop region but very long complementary ends would be more difficult to sequence both because of their length and their likelihood to form secondary structure. As DRS technology continues to improve, we hope to validate and investigate this type of cbVG species. Furthermore, the frequency at which a break-rejoin junction was sequenced by DRS didn’t always correspond with the frequency predicted by Illumina, but we cannot conclude from our data whether this is representative of the actual amount of RNA present in the sample because we did not investigate the relative efficiencies of adapter oligonucleotide binding to different RNAs.

DRS addresses the need for an affordable, timely, and sensitive tool to validate cbVG sequences. This tool is not yet quantitative, but we have seen vast improvements in the DRS kits since their initial release, which make us confident that cbVG quantitation by DRS will eventually be possible. Our sequencing outputs for cbVGs continue to increase as new kit improvements are released; to sequence cbVGs by early versions of the DRS kit, we initially had to maximize RNA concentration and supplement additional adapter oligonucleotide in the library preparation. While we sequence most of the dominant cbVG species predicted by Illumina, DRS doesn’t always capture the lower frequency cbVGs. It is also apparent that cbVG secondary structure may affect DRS outcomes; beyond physically blocking the nanopores to decrease flow cell health during DRS, cbVG translocation speeds can be variable. Longer cbVG reads that have more nucleotides available to average this translocation speed tend to be higher quality reads.

Sequence validation of cbVGs is a critical first step to study the presence of or functional impacts of molecular motifs in cbVGs. Following the recommendations herein, virologists will now be able to use DRS to sequence full-length cbVGs in their samples and use LoCA to detect and extract those reads for further study. DRS is a critical supplement to existing tools in this field, both for cbVG specialists to uncover hidden sequence features of cbVGs, and for non-specialists to discover dominant cbVGs with prospective significance in their systems.

## Materials and Methods

### Cell lines, virus infection

A549 cells were cultured and infected with SeV as previously described in (33).

### rSeVA and rSeVB virus rescue and cbVG high stocks development

Two independent rSeVC^eGFP^ viruses, named rSeVA and rSeVB, were rescued as previously described (33) and tittered as described in https://dx.doi.org/10.17504/protocols.io.ewov1d73yvr2/v1. cbVG high virus stocks were developed from the rescued viruses as described in https://dx.doi.org/10.17504/protocols.io.j8nlk9xqwv5r/v1.

### RNA extraction

Total RNA was extracted from the either the cellular or media fraction of infected cells as described in https://dx.doi.org/10.17504/protocols.io.x54v922pml3e/v1/. Cellular fractions were extracted in 1mL TRIzol reagent while media fractions (1mL) and virus stocks (500μL) were extracted in Trizol-LS reagent.

### cbVG RT-PCR and PCR bands sequencing

SeV cbVG RT-PCR was done as described in https://dx.doi.org/10.17504/protocols.io.5qpvok3j7l4o/v1. cbVG bands were cut and inserted into the pGEM-T vector as described in https://dx.doi.org/10.17504/protocols.io.yxmvme389g3p/v1. The plasmids containing cbVG bands were then sequenced through Plasmidsaurus.

### RNA sequencing, RNA-seq data preprocessing, and VODKA2 cbVG detection

RNA sequencing library preparation was performed using Illumina TruSeq Stranded Total RNA kits and sequenced at 30 million reads per sample by RNA Seq NextSeq High output 2×150. As used to generate part of Figure 5, RNA sequencing, RNA sequencing data preprocessing, and cbVG detection with VODKA2, were performed as described in (30).

### Direct RNA Sequencing library preparation

DRS libraries were prepared using sequencing kit SQK-RNA004, following the manufacturer’s instructions of the protocol and performing the optional RT step using SR+SRT-III (version DSS_9197_v4_revA_20Sep2023-minion-2). The only modification we made to this protocol is that unless otherwise specified, we designed our DRS adapter oligonucleotide to bind 32nt of the SeV trailer, instead of only the recommended 10nt (Table 1). SeV libraries were prepared using 1000ng total RNA unless otherwise indicated, while rSeV libraries were prepared using 50ng total RNA (Quantitation performed using a Qubit RNA HS assay kit on a Qubit fluorometer). All DRS experiments were sequenced for 72 hours on a MinION Mk1b using a new, previously unused FLO-MIN004RA, flow cell. Raw datasets were basecalled in MinKNOW at the high accuracy basecalling setting before passed reads were mapped to the human genome (GRCh38) using BLASTn. Nonhost reads were extracted and input into the LoCA script as described below.

**Table 1:**
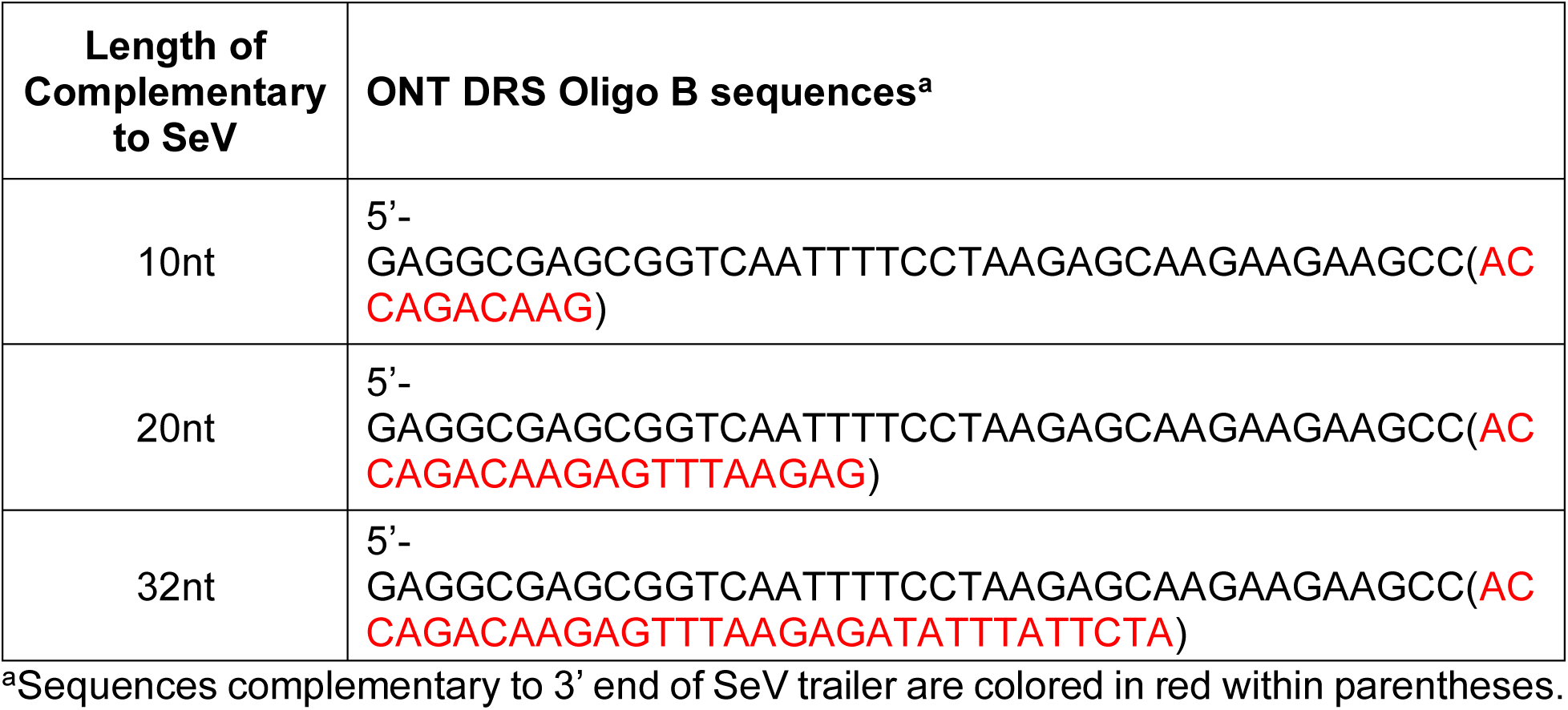
SeV DRS Adapter Oligonucleotides.

### cbVG detection with LoCA

LoCA is freely available in Github (https://github.com/lopezlab-washu/LOCA.git). BLAST nucleotide databases for the full-length virus reference genome sequences were created using BLAST 2.2.31--pl526h3066fca_3. NonHost DRS reads were input into the LoCA R script, which uses BLAST alignment and samtools to generate several files for each the virus and cbVG reads: txt file with IDs of aligned reads, fastq and fasta files of classified reads, and a LoCA report. The LoCA report summarizes the cbVG break-rejoin junctions, their predicted lengths, the size of the loop compared to the stem and the read sequence, and whether the cbVG follows the rule of 6 (can be modified or removed for viruses that are not constrained by this rule). Furthermore, LoCA outputs a mode column that is helpful for quantitating cbVGs, as it classifies into common species those cbVGs with lengths and break-rejoin junctions separated by only +/-5 nt.

### Alignment of cbVG reads

Alignment of cbVG reads for secondary analysis was performed in MegAlign software (DNAStar, Lasergene 17) using the MUSCLE algorithm. Comparisons of sequence identity were performed using the pairwise alignment function in MegAlign to align the consensus sequence from all cbVGs sharing a break-rejoin junction to a reference cbVG sharing the same break-rejoin position.

## Dava availability

Raw sequencing data of Sendai virus infections used herein are available in SRA under accession number PRJNA1232176.

## Appendices

Supplemental material is available for this article.

## Supporting information

All supplementary figures

## Acknowledgements

This work was supported by the NIH/NIAID AI137062 and funds from the BJC Investigator Program to CLB.

