## Supplementary material for "Validation of diverse and previously untraceable Sendai virus copyback viral genomes by Direct RNA Sequencing": All supplementary figures

**SUPPLEMENTAL MATERIALS**

**S1: Characterization of DRS cbVG 546 reads.** DRS output from cellular RNA extracted from A549 cells 24 hours after infection by SeV at MOI 1.5. Library was prepared using 8μM of 32bp trailer-specific oligo and sequenced for 72 hours. **(A)** Pairwise alignment of the consensus sequence of cbVG 546, obtained by aligning all SeV cbVG reads (top), to a reference cbVG 546 sequence (bottom). Mismatches in the DRS sequence are indicated as vertical grey lines against the green background. **(B)** Detailed pairwise alignment of DRS-generated cbVG 546 sequence (top) and reference cbVG546 sequence (bottom). Deletions and insertions are highlighted in grey while mismatches are highlighted in yellow. The break-rejoin junction of cbVG546 is indicated by the bold vertical red line.

**S2: Overview report for all cbVGs validated by DRS from rSeVA**. Chart shows the validated break-rejoin junction and predicted complete cbVG length, along with the number of reads by DRS, and whether it follows the paramyxovirus "rule of six" (genome length = 6n+0). Also reported are the percent of the complete cbVGs sequence validated by DRS, the percent identity of the sequenced portion of the cbVG, and a visual overview of the pairwise alignment between DRS consensus sequence and predicted reference sequence. Mismatched or un-sequenced bases are shown in grey.

**S3: Overview report for all cbVGs validated by DRS from rSeVB**. Chart shows the validated break-rejoin junction and predicted complete cbVG length, along with the number of reads by DRS, and whether it follows the paramyxovirus "rule of six" (genome length = 6n+0). Also reported are the percent of the complete cbVGs sequence validated by DRS, the percent identity of the sequenced portion of the cbVG, and a visual overview of the pairwise alignment between DRS consensus sequence and predicted reference sequence. Mismatched or un-sequenced bases are shown in grey.

**S4: Characterization of DRS host and background reads.** DRS output from cellular RNA extracted from A549 cells 24 hours after infection by SeV at MOI 1.5. Library was prepared using 8μM of 32bp trailer-specific oligo and sequenced for 72 hours. **(A)** DRS output consists of a mixture of host aligned and non-host aligned reads (percentages of total reads); the non-host reads can be further classified either as copyback viral genome (cbVG) reads, standard viral genome reads (virus), or non-host, non-virus aligned reads (background) (percentages of non-host reads). **(B)** BLASTn classification of host reads from the sample in panel A. **(C)** BLASTn classification of the Background (NonHost-NonViral) reads from panel A. **(D)** The percentage of host reads sequenced in a representative DRS experiment from Sendai virus infected LLCMK2 (black, left) and A549 (grey, right) cells.

**Fig. S1**

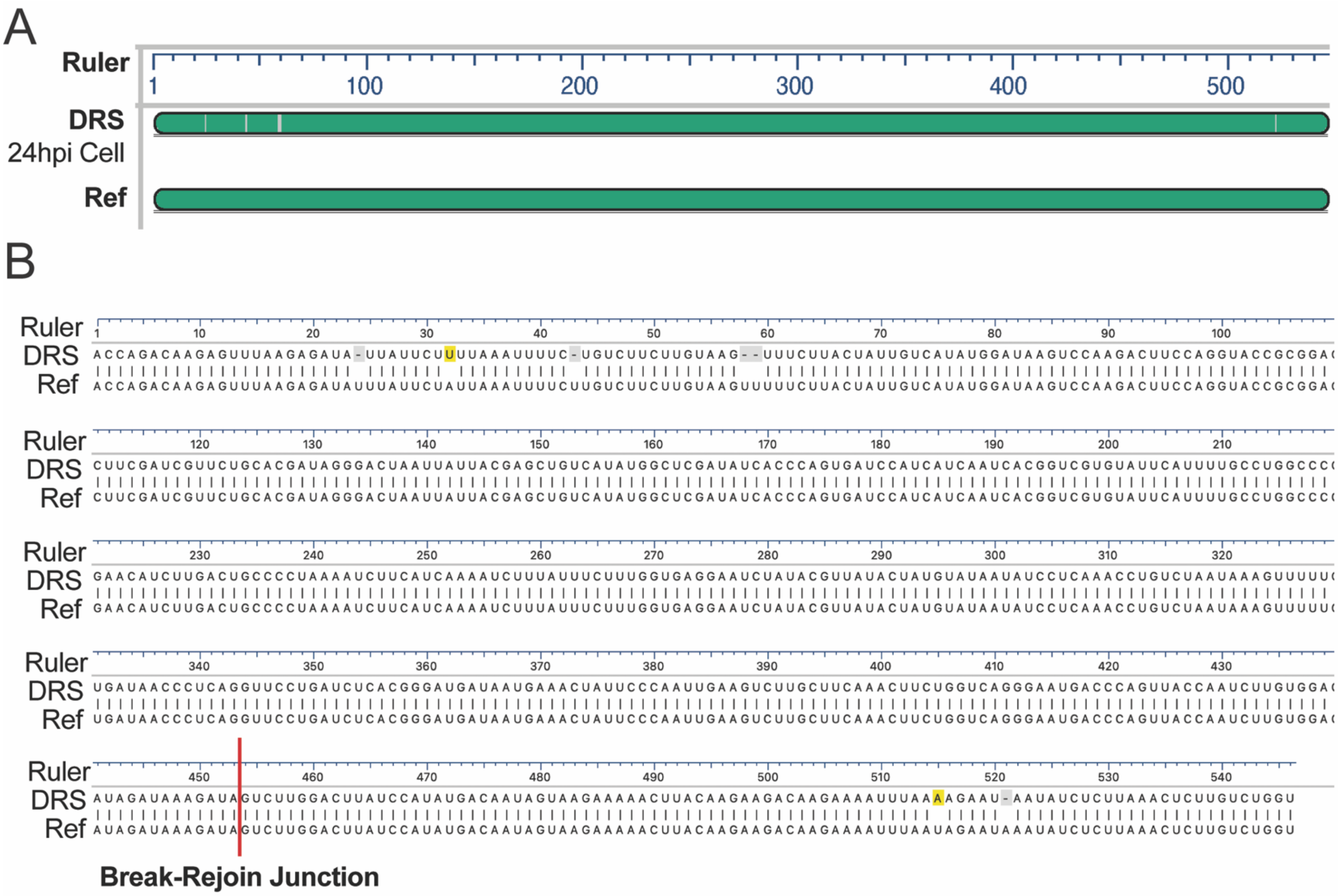

**Fig. S2**

| SeV Stock A cbVGs Sequenced by DRS | | | | | | | |
|---|---|---|---|---|---|---|---|
| **NGS Ranking** | **Break_Rejoin** | **Predicted Length** | **DRS Reads** | **Rule of Six** | **% Sequence Validated** | **% Identity** | **Alignment Overview of DRS and Reference Sequence** |
| 1 | 15,094_16,178 | 1284 | 434 | Yes | 100.0 | 99.5 | 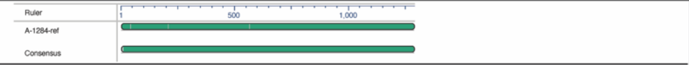 |
| 2 | 13,325_16,155 | 3078 | 182 | Yes | 100.0 | 99.6 | 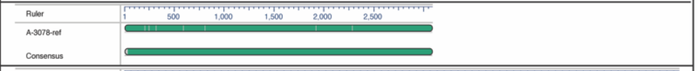 |
| 3 | 15,585_16,181 | 792 | 47 | Yes | 87.1 | 99.1 | 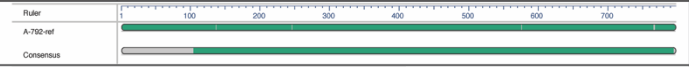 |
| 4 | 13,803_15,518 | 3240 | 227 | Yes | 100.0 | 99.5 | 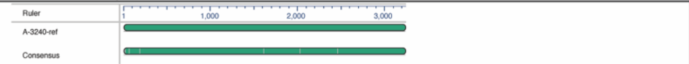 |
| 5 | 15,839_16,143 | 576 | 300 | Yes | 92.0 | 80.5 | 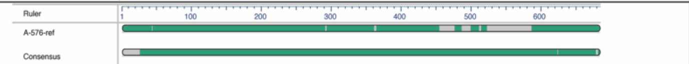 |
| 6 | 15,839_16,197 | 522 | 61 | Yes | 98.9 | 57.3 | 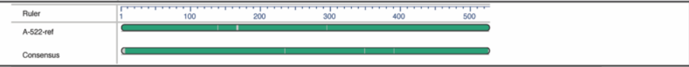 |
| 7 | 15,694_16,186 | 678 | 13 | Yes | 96.6 | 98.3 | 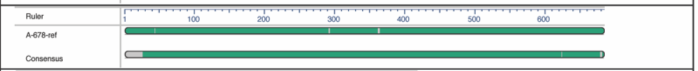 |
| 8 | 14,819_15,633 | 2106 | 21 | Yes | 100.0 | 98.9 | 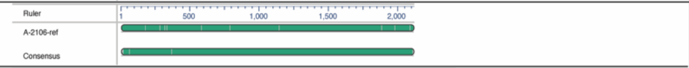 |
| 9 | 15,937_16,147 | 474 | 42 | Yes | 98.5 | 72.3 | 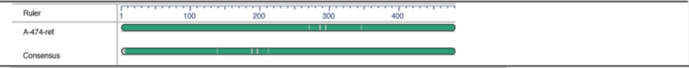 |
| 10 | 15,389_16,084 | 1086 | 14 | Yes | 100.0 | 97.9 | 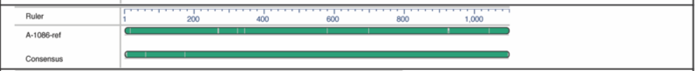 |
| 11 | 16,058_16,104 | 396 | 479 | Yes | 100.0 | 93.0 | 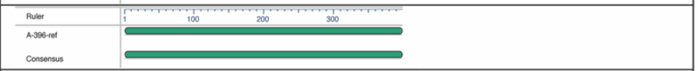 |
| 13 | 15,200_16,103 | 1254 | 17 | Yes | 100.0 | 97.7 | 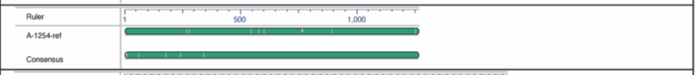 |
| 14 | 15,506_15,960 | 1092 | 3 | Yes | 33.0 | 96.0 | 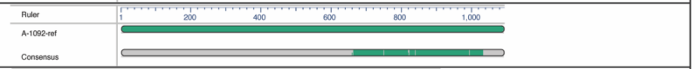 |
| 15 | 15,092_15,655 | 1806 | 3 | Yes | 74.0 | 72.6 | 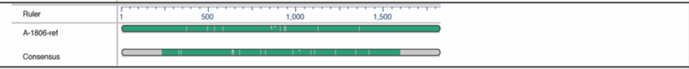 |
| 17 | 1_15,727 | 16,830 | 3 | Yes | 10.8 | 97.4 | 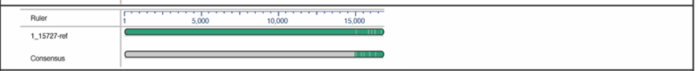 |
| 18 | 16,055_16,095 | 408 | 45 | Yes | 95.3 | 88.9 | 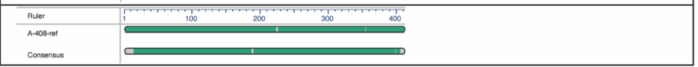 |

**Fig. S3**

| SeV Stock B cbVGs Sequenced by DRS | | | | | | | |
|---|---|---|---|---|---|---|---|
| **NGS Ranking** | **Break_Rejoin** | **Predicted Length** | **DRS Reads** | **Rule of Six** | **% Sequence Validated** | **% Identity** | **Alignment Overview of DRS and Reference Sequence** |
| **1** | 14,822_16,188 | 1548 | 1058 | Yes | 100.0 | 99.8 | 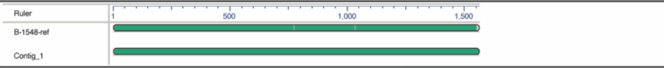 |
| **2** | 12,858_16,016 | 3684 | 25 | Yes | 95.4 | 99.0 | 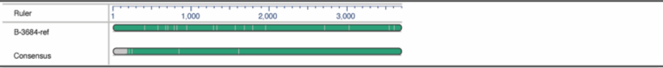 |
| **3** | 15,660_16,130 | 768 | 44 | Yes | 100.0 | 99.0 | 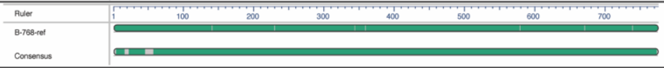 |
| **4** | 12,238_14,926 | 5394 | 3 | Yes | 41.5 | 95.5 | 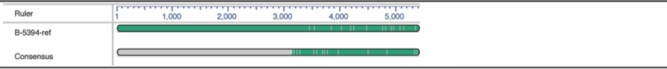 |
| **5** | 15,416_15,690 | 1452 | 10 | Yes | 90.1 | 86.6 | 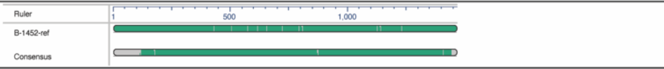 |
| **6** | 15,315_15,965 | 1278 | 5 | Yes | 86.5 | 96.9 | 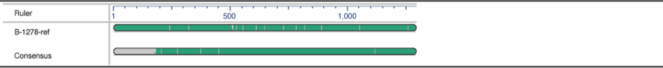 |
| **7** | 15,569_16,101 | 888 | 4 | Yes | 73.6 | 98.3 | 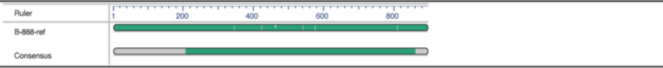 |
| **9** | 1_15,745 | 16,812 | 4 | Yes | 12.9 | 97.3 | 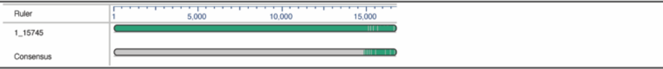 |
| **12** | 5,738_16,174 | 10,656 | 3 | Yes | 56.0 | 95.6 | 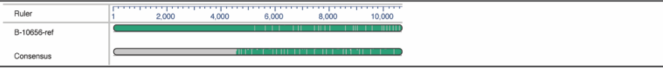 |
| **133** | 16,058_16,074 | 426 | 4 | Yes | 50.0 | 96.2 | 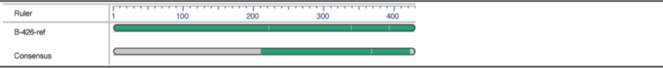 |

Fig. S4

A

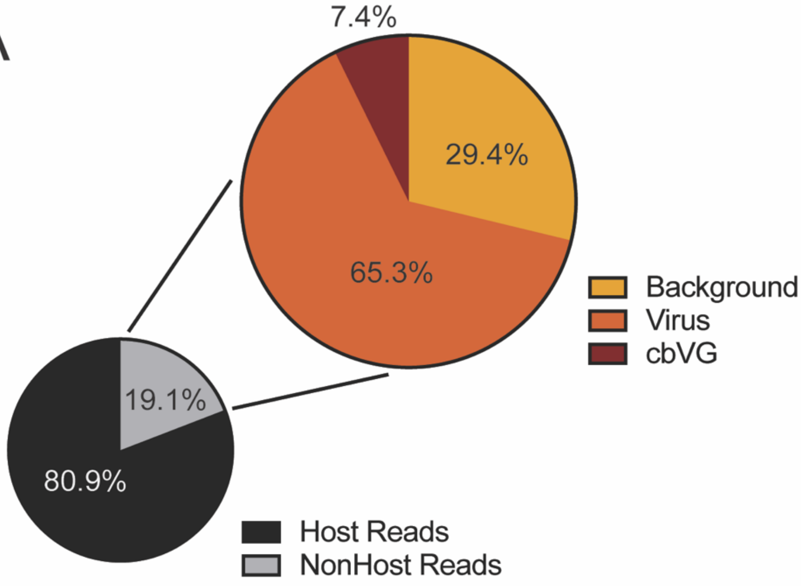


B

**Host Reads**

| Reads | BLASTn Classification |
|---|---|
| 965 | Chromosome 17 |
| 240 | 28S Ribosomal RNA |
| 172 | 18S rRNA |
| 104 | lncRNA |
| 29 | Mitochondrion |
| 16 | Other |

C

**Background Reads**

| Species | BLASTn Classification |
|---|---|
| 1 | No classification, GC-repeats |
| 2 | No classification, AU-repeats |
| 3 | Chromosome 17 |
| 4 | lncRNA |

D

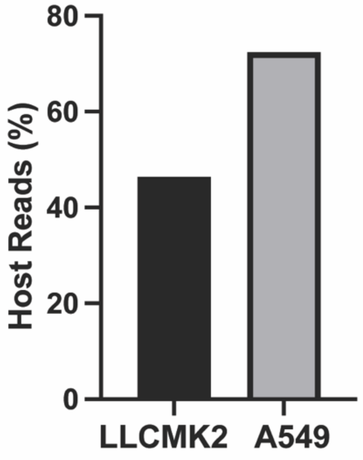
